# Reproductive functions and genetic architecture of the seminal fluid and sperm proteomes of the mosquito *Aedes aegypti*

**DOI:** 10.1101/405431

**Authors:** Ethan C. Degner, Yasir H. Ahmed-Braimah, Kiril Borziak, Mariana F. Wolfner, Laura C. Harrington, Steve Dorus

**Affiliations:** Department of Entomology, Cornell University, Ithaca, NY; Department of Molecular Biology and Genetics, Cornell University, Ithaca, NY; Center for Reproductive Evolution, Syracuse University, Syracuse, NY

## Abstract

The yellow fever mosquito, *Aedes aegypti,* transmits several viruses, including dengue, Zika, and chikungunya. Some proposed efforts to control this vector involve manipulating reproduction to suppress wild populations or replacing them with disease-resistant mosquitoes. The design of such strategies requires an intimate knowledge of reproductive processes, yet our basic understanding of reproductive genetics in this vector remains largely incomplete. To accelerate future investigations, we have comprehensively catalogued sperm and seminal fluid proteins (SFPs) transferred to females in the ejaculate using tandem mass spectrometry. By excluding female-derived proteins using an isotopic labelling approach, we identified 870 sperm proteins and 280 seminal fluid proteins. Functional composition analysis revealed parallels with known aspects of sperm biology and SFP function in other insects. To corroborate our proteome characterization, we also generated transcriptomes for testes and the male accessory glands—the primary contributors to *Ae. aegypti* sperm and seminal fluid, respectively. Differential gene expression of accessory glands from virgin and mated males suggests that protein translation is upregulated post-mating. Several SFP transcripts were also modulated after mating, but >90% remained unchanged. Finally, a significant enrichment of SFPs was observed on chromosome 1, which harbors the male sex determining locus in this species. Our study provides a comprehensive proteomic and transcriptomic characterization of ejaculate production and composition and thus provides a foundation for future investigations of *Ae. aegypti* reproductive biology, from functional analysis of individual proteins to broader examination of reproductive processes.

## Introduction

The mosquito, Aedes aegypti, is the most important vector of arboviruses globally, transmitting viruses that cause dengue [1], Zika [2], chikungunya [3], and yellow fever [4]. Consequently, Ae. *aegypti* places a severe strain on public health infrastructure around the world [5]. Despite decades of effort to control mosquito populations, *Ae. Aegypti* continues to drive human epidemics of these diseases. New and improved control strategies are needed to prevent future outbreaks and mitigate disease burden.

Some promising control strategies under development target reproduction to suppress mosquito populations. For example, sterilized males can be released to suppress populations by impairing reproduction by their wild mates [6, 7]. Manipulating reproductive phenotypes may also provide a means of driving disease-refractory traits into a population (reviewed in [8]). One such strategy employs the intracellular bacterium Wolbachia, which, when introduced into *Ae. aegypti,* induces cytoplasmic incompatibility that allows the bacterium to spread in a population, potentially to fixation [9]. Cytoplasmic incompatibility causes sperm of males with Wolbachia to be incompatible with uninfected females’ eggs, whereas Wolbachia-positive females can reproduce with any male, regardless of infection status (reviewed in [10]), giving Wolbachia-positive individuals a fitness advantage over their uninfected counterparts. This bacterium also blocks or reduces transmission of several viruses, including dengue [11] and Zika [12]. Consequently, introduction of novel Wolbachia infections into vector populations is being explored as a transmission reducing strategy.

Designing mosquito control strategies that target reproduction requires an intimate knowledge of the cellular and molecular mechanisms that govern it. Yet, only a few functions of proteins involved in mosquito reproduction have been described to date. For example, in *Ae. aegypti* seminal fluid proteins (SFPs) induce several physiological and behavioral changes in females, including refractoriness to future mating [13-18], stimulation of oogenesis [19], enhanced survival [18], and the ability to fertilize eggs [20]. However, the molecular identity of active SFP components for this species remains elusive. Seminal fluid initiates sperm motility via the action of proteases in many insects (silkworm [21]; water strider [22]; Culex mosquito [23]), but the precise sperm proteins on which seminal fluid acts in *Ae. aegypti* have not been identified. Similarly, sperm-associated odorant receptors may control motility in *Ae. aegypti,* although the exact function and ligands of these receptors are unknown [24]. Finally, the mechanism by which Wolbachia induces cytoplasmic incompatibility has not been described in *Ae. aegypti,* but Wolbachia proteins contained in sperm are hypothesized to be involved [25,26].

Identification of sperm proteins and SFPs that are transferred to females during copulation is an important objective to enable future investigations into specific reproductive processes. Components of the transferred ejaculate include sperm and seminal fluid—both of which play vital roles in mosquito reproduction. An *Ae. aegypti* seminal fluid proteome was first reported by Sirot *et al.* [27] based on mass spectrometry analyses. That study described 93 putative SFPs transferred during mating. While not a primary focus of their work, they also identified 101 putative sperm proteins. Later work identified more than twice as many SFPs from *Ae. albopictus* using a similar method [28]. Proteome complexity of other insects’ seminal fluid (reviewed in [29-31]) and sperm [32-34] suggests that more proteins remain to be identified in the *Ae. aegypti* ejaculate.

Here, we build on the foundational work of Sirot *et al.* [27], using tandem mass spectrometry (MS/MS) with greater sensitivity to identify the constituent proteins of both *Ae. aegypti* sperm and seminal fluid. We also profile the transcriptomes of the male accessory glands (MAG; before and after mating) and testes, the major source tissues for SFPs and sperm, respectively. Our proteomic characterization represents a nearly four-fold expansion of putative SFPs and a more than eight-fold expansion in the *Ae. aegypti* sperm proteome. Our results yield insights into the molecular function, genome organization, regulation, and evolution of sperm proteins and SFPs in this important disease vector. Ultimately, these proteomes provide a basis for future studies of mosquito reproduction and, potentially, a catalog of molecular targets for the development of novel mosquito control methods.

## Materials and Methods

### Rearing

Mosquitoes were derived from a laboratory colony of *Ae. aegypti* that was established from individuals collected in Bangkok, Thailand, in 2011 and supplemented with wild caught mosquitoes every 2 - 3 y. Mosquitoes were reared as described previously by Degner and Harrington [35]. Briefly, eggs were hatched under vacuum pressure, and a day later 200 first instar larvae were transferred to rearing trays of 1 L deionized water with four fish food pellets as diet (except in the case of ^15^N-labeled females; see below). Pupae were separated by sex based on size, allowed to eclose individually in separate test tubes, and adults were transferred into single-sex cages with 10% sucrose provided *ad libitum.*

### Sperm isolation

Males used for sperm protein sample preparation were aged between 5 and 8 days post-eclosion (dpe). Mature sperm were isolated from the seminal vesicles of each male. Males were dissected in physiological saline (133 mM NaCl, 2.63 mM KCl, 9.75 mM Na_2_HPO_4_, 3 mM KH_2_PO_4_, 2 mM CaCl_2_, adjusted to pH 6.9; hereafter “saline”). The seminal vesicle was isolated from other tissue and consecutively transferred to two clean droplets of saline to remove any adherent fat body or other debris. Clean dissecting tools were used to transfer the seminal vesicle to a final droplet of saline where it was ruptured to release sperm. Sperm suspended in saline were transferred to a microcentrifuge tube on ice, and pooled sperm samples were flash frozen in liquid nitrogen every 2 h.

Two biological replicates included sperm combined from 400 and 470 randomly selected males, respectively. Pooled samples were centrifuged at 25,000 x g and the supernatant was removed, leaving 18μl saline with the pellet. An equal volume of 2x Laemmli buffer + 5%,0-mercaptoethanol was added to the pellet, and samples were solubilized by sonicating for 30 s, boiling for 15 min, and re-sonicating for 30 s. To remove any particulate debris, samples were spun down at 10,000 x g for 10 min, and the supernatant was placed in a fresh tube. Protein was quantitated using a 1:169 5 dilution of the sample using the EZQ assay (Thermo Fisher Scientific, Waltham, MA). Protein used for subsequent mass spectrometry was standardized across biological replicates (16μg).

### Transferred ejaculate esolation

Males were reared as described above. As larvae, females were labelled with ^15^N using the rearing methodology of Sirot *et al.* [27]. Briefly, a prototrophic yeast strain (D273-10B) was grown in media whose only nitrogen source was ^15^N ammonium sulfate (Cambridge Isotope Laboratories; Cambridge, MA). Yeast were grown to saturation, pelleted, and resuspended in PBS to a final volume of one sixteenth of the growth media. A few drops of yeast slurry were provided to newly hatched first instar larvae after vacuum hatching. One day after hatching, 200 larvae were placed in a rearing tray with 200 mL water from a previous cohort of ^15^N-yeast-reared mosquitoes and 800 mL of deionized water. Larvae were fed 4 mL of labeled yeast slurry a day after hatching, and again at 4 d after hatching. Pupae were isolated in individual tubes to ensure virginity upon eclosion, and females were put in an 8 L bucket cage with sucrose solution ad libitum. Sucrose was replaced every 2 d to preclude the introduction of unlabeled nitrogen via microbial contamination.

At 4 - 5 dpe, matings between labeled females and unlabeled males were observed as in Degner and Harrington [35]. After mating, females were immediately placed on ice and dissected within 3 min. Because mosquito seminal fluid is known to contain proteases [27], mosquitoes were dissected in saline with protease inhibitors (cOmplete Mini Protease Inhibitor Cocktail; Sigma Aldrich, St. Louis, MO). In contrast to Sirot *et al.* [27], we dissected only bursae (and not spermathecae), cutting the bursae just proximal to the vaginal lips. Upon excision, bursae (n = 35 - 43 per replicate) were transferred to 33 L 1x Laemmli buffer diluted from a 2x stock solution with saline, -mercaptoethanol and 1x protease inhibitors. Samples were sonicated for 30 s, boiled for 10 min, sonicated for 30 s, and centrifuged at 10,000 x g for 10 min. The supernatant was removed; 30 L were frozen at -80°C, and 2μL were diluted 1:3 with buffer for protein quantitation using the EZQ assay (Thermo 194 Fisher Scientific, Waltham, MA). In parallel, we also prepared samples of bursae from virgin, labeled females from each cohort to assess the efficiency of ^15^N-labeling. Mated and virgin bursae samples contained 18 and 8 g of protein, respectively.

### Tandem mass spectrometry analysis

Solubilized proteins were separated on a 1-dimensional SDS-PAGE gel and split into 6 fractions, with two biological replicates run in parallel (Figure S1). Gel fractions were reduced in dithiothreitol, alkylated with iodoacetamide, and digested with trypsin. Lyophilized, digested proteins were reconstituted in 0.5% formic acid and subjected to nanoLC-ESI-MS/MS analysis using an Orbitrap Fusion Tribrid mass spectrometer (Thermo-Fisher Scientific, San Jose, CA) equipped with nanospray Flex Ion Source, and coupled with a Dionex UltiMate 3000RSLC nano system (Thermo, Sunnyvale, CA). Peptide samples were injected onto a PepMap C-18 RP nano trap column (5μm, 100μm i.d. x 20mm, Dionex) with nanoViper fittings at 20μL/min flow rate for desalting. Samples were then separated on a PepMap C-18 RP nano column (2μm, 75μm x 15cm) at 35°C, followed by elution on a 90 min gradient of 5% to 35% acetonitrile in 0.1% formic acid at 300 nL/min. Finally, a 5 min ramping to 90% acetonitrile in 0.1% formic acid and a 5 min hold at this eluent completed each run cycle. Between cycles, the column was re-equilibrated for 25 min using 0.1% formic acid. The Orbitrap Fusion was run in positive spray ion mode with spray voltage set at 1.6 kV and a source temperature at 275°C. External calibration for FT, IT, and quadrupole mass analyzers was performed. In data-dependent acquisition analysis, the instrument was operated using FT mass analyzer in MS scan to select precursor ions followed by 3 s “Top Speed” data-dependent CID ion trap MS/MS scans at 1.6 m/z quadrupole isolation for precursor peptides with multiple charged ions above a threshold ion count of 10,000 and normalized collision energy of 30%. MS survey scans at a resolving power of 120,000 (fwhm at m/z 200), for the mass range of m/z 375-1575. Dynamic exclusion parameters were set at repeat count 1 with a 20 s repeat duration, an exclusion list size of 500, and 40 s of exclusion duration with ±10 ppm exclusion mass width. The activation time was 10 ms 219 for CID analysis. All data were acquired under Xcalibur 3.0 operation software (Thermo-Fisher Scientific). All post-quantitation sample preparation was conducted at the Cornell Biotechnology Resource Center.

### Peptide identification and protein annotation

Raw data from each MS/MS run was analyzed by X!Tandem [36] and Comet [37] against the *Ae. aegypti* L5.0 protein database (GCF_002204515.2; [38]). Only the longest protein isoform of each gene was included in the search database, resulting in a database of 14,626 proteins. For X!Tandem, a fragment ion mass tolerance of 0.40 Da and a parent ion tolerance of 10.0 PPM were used. For Comet, a fragment bin tolerance of 1.0005 with a 0.4 offset and a parent ion tolerance of 10.0 PPM were used. Iodoacetamide derivative of cysteine was specified as a fixed modification, whereas oxidation of methionine and deamidation of glutamine and asparagine were specified as variable modifications. Peptides were allowed up to two missed trypsin cleavage sites. All downstream analyses were conducted using the Trans-Proteomic Pipeline (TPP v5.0 (Typhoon) rev 0; [39]). False Discovery Rates (FDRs) were estimated with a randomized decoy database using PeptideProphet [40], employing accurate mass binning model and the nonparametric negative distribution model. X!Tandem and Comet PeptideProphet results were merged using iProphet [41], to provide more robust peptide identification. Peptide identifications were accepted if they could be established at greater than 95.0% iProphet probability, and protein assignations were accepted if they could be established at greater than 99.0% probability. Proteins that contained identical peptides and could not be differentiated based on MS/MS analysis alone were grouped to satisfy parsimony principles.

### Verification of labelling efficiency

For ejaculate samples, protein from virgin female controls was run on the same gel as protein from their mated counterparts, and labeling efficiency was verified on a representative fraction in each cohort’s virgin sample (Figure S1). No peptides or proteins were identified using our statistical criteria in the labelled virgin female samples when searched using standard, unlabelled mass parameters. Thus, whole female labelling was complete in relation to MS/MS peptide and protein identification and precluded the identification of female proteins from mated female bursae.

### Protein quantitation

Protein quantitation was conducted using the semi-quantitative spectral counting approach implemented by the APEX Quantitative Proteomics Tool [42]. The top 50 identified proteins with the highest protein identification probabilities were utilized as the training dataset. The 35 physicochemical properties available in the APEX tool were used for prediction of peptide detection/non-detection in the construction of a training dataset file. Protein probabilities (O_*i*_) were computed using the Random Forest classifier algorithm trained with the data set generated in the previous step. APEX protein abundances per sample were calculated using the protXML file generated by ProteinProphet.

### Experimental design and statistical rationale

To ensure the reproducibility of protein identifications, two biological replicates of each tissue were analyzed and robust identification criteria were applied (see above). Each replicate was prepared from independent cohorts of mosquitoes. To control for the possibility that unlabelled female-derived proteins were identified in our ejaculate samples, we assessed labelling efficiency by conducting mass spectrometry on virgin bursae alone, using a representative gel slice from each biological replicate (Figure S1). False Discovery Rates (FDRs) were estimated with a randomized decoy database using PeptideProphet [40], employing accurate mass binning model and the nonparametric negative distribution model. Corrections for multiple testing were applied where appropriate to ensure the conservative nature of statistical tests.

### Transcriptome analysis of testes and male accessory glands

Testes were harvested from males at 1 dpe and transferred to TRIzol. Because mature sperm are actively produced at this age [43], and spermatogenesis is at its peak [44,45], testes at this age likely contain the majority of transcripts that contribute to the testicular sperm proteome. Male accessory glands (MAG) including the connecting ejaculatory duct were dissected from virgin males aged 6 and 8 dpe. We also analyzed MAG from mated males at the same age. Previous work has demonstrated that *Ae. aegypti* males become depleted after mating with three to five females in succession, and seminal fluid is slowly regenerated over 48 h [46]. In our study, we provided males with four virgin females for a period of 8 h to allow for male seminal fluid depletion. On average, each male mated with more than three females in this period (as determined by dissection of females’ spermathecae). Males were dissected in saline 16 h after their female mates had been removed. We generated four biological replicates from independent cohorts for each treatment, and each replicate contained combined tissue from 20-40 (testes) or 40-60 (MAG) males. Total RNA was extracted from each sample in Trizol following manufacturer’s instructions (Invitrogen, Carlsbad, CA). Poly-A mRNA was isolated and cDNA libraries were prepared using the QuantSeq 3’ mRNA-Seq Library Prep Kit FWD for Illumina (Lexogen, Vienna, Austria). Amplified cDNA products were run on an AATI Fragment Analyzer (Advanced Analytical Technologies, Inc.; Hialeah, FL) so that the cDNA was of sufficient concentration for sequencing. Library concentrations were balanced using digital PCR [47], and each of the 12 uniquely barcoded samples were sequenced in one lane using the Illumina HiSeq 2500 platform with 100 bp reads. All sequencing was conducted at the Cornell Biotechnology Resources Center.

For further analysis of the transcriptomes, we included additional, publicly available data to evaluate tissue-biased gene expression. These include gonadectomized male carcass (SRP075464; [48]) and a virgin female reproductive tract sample (SRP068996; [49]). Raw RNA-seq reads were processed by trimming the first 10 bases from the 3’ position, followed by quality trimming of both ends to a minimum quality Phred score of 20 (Sickle v.1.210; [50]). Processed reads were 293 then mapped to the *Ae. aegypti* genome (VectorBase release L5.1; [38]) using Hisat2 (v.2.1.0; [51]) with default parameters, and transcript abundance was estimated for each sample with StringTie (v.1.3.4; [52]). Raw counts for each sample were extracted from the StringTie abundance estimates using the auxiliary “prepDE.py” script provided on the StringTie website (https://ccb.jhu.edu/software/stringtie/). Signal peptides in the translated transcriptome were predicted *in silico* using a local installation of SignalP (v.4.1; [53]).

We used raw counts from the RNA-seq samples to (1) classify genes based on tissue-biased expression in the MAG and testes, and (2) identify genes differentially expressed between virgin and mated MAG. Count matrices were filtered to remove low abundance transcripts (counts per million < 5). First, we compared expression levels between the focal tissue (testes or virgin MAG) to three other tissues: gonadectomized male carcass [48], virgin female reproductive tract [49], and virgin MAG or testes (present study). We classified genes as testes- or MAG-biased if they had >2-fold higher transcript abundance compared to other samples at a minimum FDR cutoff of 0.05 (edgeR v.3.23.2; [54]). Finally, we identified differentially expressed genes between virgin and mated MAGs as having >2-fold abundance difference at an FDR cutoff of 0.01. Using the same differential expression criteria, we also re-analyzed data from female reproductive tracts of virgin and just-mated females [49] to identify transcripts putatively transferred to females in the ejaculate using current annotated gene models.

### Chromosomal distribution of male reproductive genes

To evaluate the chromosomal distribution of SFPs, sperm proteins and MAG/testes-biased genes, we calculated the expected number of genes for each class on each chromosome, assuming a random distribution of genes across the genome. We then multiplied the total number of genes within each class by the expected proportion for each chromosome (based on the proportion of total genes on that chromosome) to establish an observed/expected ratio. We also calculated this ratio for a 123 Mb region on chromosome 1 that surrounds the sex determining locus and has low rates of 317 recombination [55]. A 318 x^2^ test *(d.f.* = 1) for each gene class per chromosome was used to test for biased representation.

### Orthology relationships and gene ontology analysis

Species orthology was assessed using a local installation of OrthoDB with default BLAST and clustering parameters [56]. Protein sequences for *Ae. aegypti* and *Ae. albopictus* were retrieved from NCBI (GCF_002204515.2 and GCF_001876365.2, respectively), and *Drosophila melanogaster* protein sequences were retrieved from FlyBase (r6.18). Protein sequences for each species were filtered to retain only the longest isoform for each gene. Sperm proteins and SFPs for *Ae. albopictus* and *D. melanogaster* were based on previous mass spectrometry based proteomic studies [28,32,57]. Because SFP identifications by Boes *et al.* [28] were made using a *de novo* transcriptome (the *Ae. albopictus* genome had not yet been released), we converted the original SFP identifications to their current accession numbers by blasting assembled transcripts from Boes *et al.* [28] to the assembled *Ae. albopictus* genome [58]. All Gene Ontology (GO) analyses were conducted using GOseq [59], with gene lengths derived from the longest transcript of each assembled StringTie gene. GO terms were extracted from BLAST results against the SwissProt database (www.uniprot.org).

### Data Availability

Mass spectrometry proteomics data have been deposited to the ProteomeXchange Consortium (http://proteomecentral.proteomexchange.org) via the PRIDE partner repository with the dataset identifier PXD010293 and 10.6019/PXD010293. The R analysis pipeline used for transcriptomic analysis is available as part of this project’s GitHub repository (https://github.com/YazBraimah/Aegypti.ejaculatomics). Raw RNA-seq reads from this study can be obtained from the Sequence Read Archive (SRA) accession SRP158536. RNA-seq data from Sutton *et al.* [48] and Alfonso-Parra *et al.* [49] can be accessed at SRA accessions SRP075464 and SRP068996, respectively.

## Results

### Sperm and ejaculate proteome characterization

To characterize *Ae. aegypti* proteins transferred to females during mating and distinguish between sperm and non-sperm seminal fluid components, we used MS/MS to analyze proteins from (1) purified sperm isolated from male seminal vesicles and (2) whole ejaculate from bursae of mated females. For our analysis of the whole ejaculate, females were labelled with heavy nitrogen to preclude detection of female-contributed proteins. Labelling efficiency was determined to be complete with regard to peptide and protein identification (see Materials and Methods). Two replicates were analyzed per sample. High levels of reproducibility were observed, including 85% of sperm proteins and 78% of ejaculate proteins identified in both biological replicates. Similar levels of inter-replicate consistency have been reported for sperm proteomes of other organisms [34]. To maximize protein identification, we combined spectra across biological replicates for our final proteome determination, resulting in 54,894 peptide spectral matches (PSMs) for sperm, and 30,801 for the ejaculate (Table S1). The nearly two-fold disparity in PSMs is due to the contribution of labelled female proteins in the ejaculate sample. In total, 870 and 811 proteins with at least 2 unique PSMs were identified in sperm and ejaculate, respectively (Table S2). Sperm proteins were identified by an average of 11.4 unique PSMs and 62.8 total PSMs per protein. Ejaculate proteins were identified with an average of 8.0 unique PSMs and 37.8 total PSMs. As was expected given the substantial contribution of sperm cells to ejaculate composition, extensive overlap was observed between sperm and ejaculate proteomes; 516 proteins were detected in both samples, while 354 proteins were only identified in sperm and 295 proteins were uniquely detected in the ejaculate (Figure 1A).

**Figure 1.**
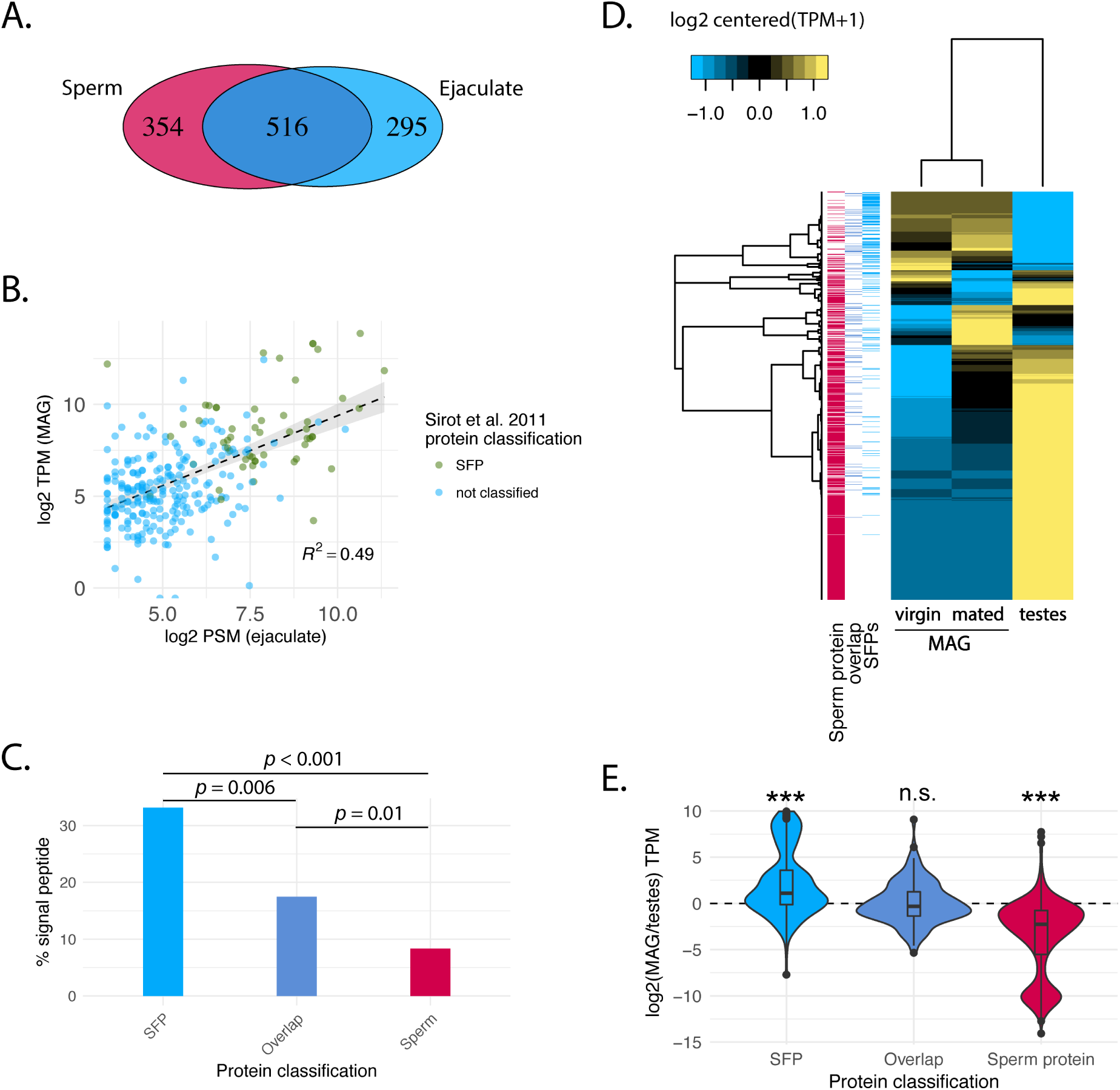
Characteristics of the *Ae. aegypti* sperm and ejaculate proteomes. **(A)** Venn diagram representing the number of proteins identified by ≥2 unique peptides using LC-MS/MS in sperm (red) and ejaculate (blue) proteomes. **(B)** Scatter plot of the protein (log2PSM) and mRNA (log2TPM) abundances of the 177 SFPs and 103 overlap SFPs. Proteins classified as SFPs in the Sirot *et al.* study are highlighted in green and those not identified in Sirot *et al.* [27] are highlighted in blue. Dotted line is a linear model fit, and gray shading represents the 95% confidence interval. **(C)** Percentage of proteins containing a predicted signal peptide sequence among SFPs, sperm, and SFP/sperm overlap protein classes; all proportions are significantly different from each other (%^2^, p < 0.05). **(D)** Heatmap representing the mRNA abundance profile of male reproductive classes identified in this study. Cladograms on the left and top represent Pearson cluster grouping of genes and samples, respectively. Annotation bars on the left indicate the protein classification for each gene. **(E)** Violin plots displaying the fold-change (log2) distribution of mRNA abundance between MAG and testes samples for each male reproductive protein class. Width of violins represents the number of proteins at any given TPM ratio; boxes represent inner quartiles and outliers drawn using the Tukey method. Asterisks indicate groups that are significantly different from each other (Wilcoxon Signed rank test; *** = *p <* 0.001).rank sum test; *W* = 3786, *p <* 0.001), consistent with the higher sensitivity and coverage of new MS/MS methods utilized in this study. Proteins identified by Sirot *et al.* [27] also exhibited significantly higher levels of MAG expression (Figure 1B; Wilcoxon rank sum test; *W* = 3005, *p <* 0.001). We therefore conclude that the sensitivity of our SFP characterization resulted in the addition of a far greater number of low abundance SFPs.

The primary goal of this study was to use MS/MS with higher sensitivity and accuracy to expand upon the prior characterization of *Ae. aegypti* SFPs and sperm by Sirot *et al.* [27]. They identified 74 SFPs that mapped to the recently refined *Ae. aegypti* genome [38]; some of the 93 SFPs described by Sirot *et al.* [27] do not map to the new genome or are now part of larger, fused gene models. Of these, we identified 60 (81%) in our ejaculate sample, 32 of which were also identified in our purified sperm sample. It is noteworthy that we detected an additional 5 SFPs from Sirot *et al.* [27], but these were not included in our SFP list because they did not meet our two unique peptide inclusion criteria. As such our proteomic characterization expands the previous *Ae. aegypti* SFP characterization.

### Refined seminal fluid protein classification

Because the ejaculate is a complex mixture of sperm and seminal fluid, we applied stringent inclusion criteria to our ejaculate samples to refine the delineation between SFPs and sperm proteins. We first removed 10 proteins involved in protein translation (*i.e.* ribosomal proteins, translation initiation factors, and elongation factors; Table S2) from the list of putative SFPs. These proteins exhibit ubiquitous patterns of expression, including both MAG and testes, and are unlikely to be bona fide secreted SFPs. Although we cannot rule out that they are secreted SFPs, their presence may be the result of cell rupture during seminal secretion [60] or transfer of MAG cells to the female, as has been described in *D. melanogaster* [61]. To reduce the possible inclusion of sperm proteins that were absent in our sperm proteome but present in the ejaculate (perhaps due to low abundance), we define “high confidence SFPs” as proteins with a minimum of 6 total PSMs in the ejaculate, 2 unique PSMs, and not present in our sperm proteome; this resulted in 177 high-confidence SFPs (Table S2).

Previous analyses of insect sperm proteomes have consistently identified proteins generally considered to be SFPs (*i.e.* highly expressed in the MAG and believed to be secreted molecules transferred to females as non-sperm components [32,62,63]. To identify proteins predominantly produced by the MAG, but identified in both our sperm and ejaculate sample, we used a 2.5-fold greater abundance threshold in the ejaculate relative to sperm. This resulted in the identification of 103 additional putative SFPs, which we label as “sperm/SFP overlap” (Table S2), including 53 with 5-fold greater protein abundance in the ejaculate relative to sperm. In total, this results in a combined SFP proteome of 280 proteins.

To better understand the relationship between candidate SFPs identified in the current study and by Sirot *et al.* [27], we examined protein abundance and transcript levels of both protein sets (50 of the 74 SFPs identified by those authors are present in our SFP proteome). This revealed that SFPs identified by Sirot *et al.* [27] were significantly more abundant than the remainder of our 280 SFPs (Figure 1B); Wilcoxon

To evaluate our SFP characterization, we explored three types of analyses. First, we determined the presence of predicted signal peptides in identified proteins — a hallmark of secreted proteins. Amongst our high-confidence SFPs, μ33% contained signal peptides, in comparison to only μ9% of sperm proteins (x^2^ = 69.2, *p <* 0.001). Additionally, ˜17% of the sperm/SFP overlap proteins had a predicted signal peptide — also significantly more than the sperm proteome (x^2^ = 6.47, *p* = 0.01), and less than high-confidence SFPs (x^2^ = 7.7, *p* = 0.006; Figure 1C). Second, we used RNA-seq to characterize the MAG and testis transcriptome, the predominant source tissues of proteins in our analysis. As expected, SFPs (Wilcoxon Signed rank test; *p <* 0.001) and sperm proteins (Wilcoxon Signed rank test; *p <* 0.001) exhibit MAG- and testis-biased expression, respectively, while the sperm/SFP overlap proteins exhibit less tissue-biased expression levels overall (Wilcoxon Signed rank test; *p* = 0.36; Figure 1D-E). Third, we assessed the amount of high-confidence SFP protein abundance variation explained by variation in MAG expression levels. This revealed a significant correlation between protein and transcript abundance (R^2^ = 0.49, *F* = 88.24, *p <* 0.001; Figure 1B). In conjunction, these results support our proteomic characterization of SFPs secreted from the MAG, including proteins present in both sperm and ejaculate samples but highly enriched in the ejaculate proteome.

### Gene ontology of sperm and seminal fluid proteomes

Gene ontology (GO) categories enriched in mosquito sperm are largely similar to those found in other insects’ sperm (Table 1, Table S3; [32,34,62,63]). Proteins associated with mitochondria and the axoneme are the most enriched cellular components in the sperm proteome. Proteasome components are also enriched. Over-represented biological processes in sperm include nucleotide biosynthesis, metabolic processes related to the tricarboxylic acid cycle, and proteins regulating ciliar function. Nucleotide binding, ion transport, and oxidoreductase activity are enriched molecular functions in sperm.

**Table 1.**
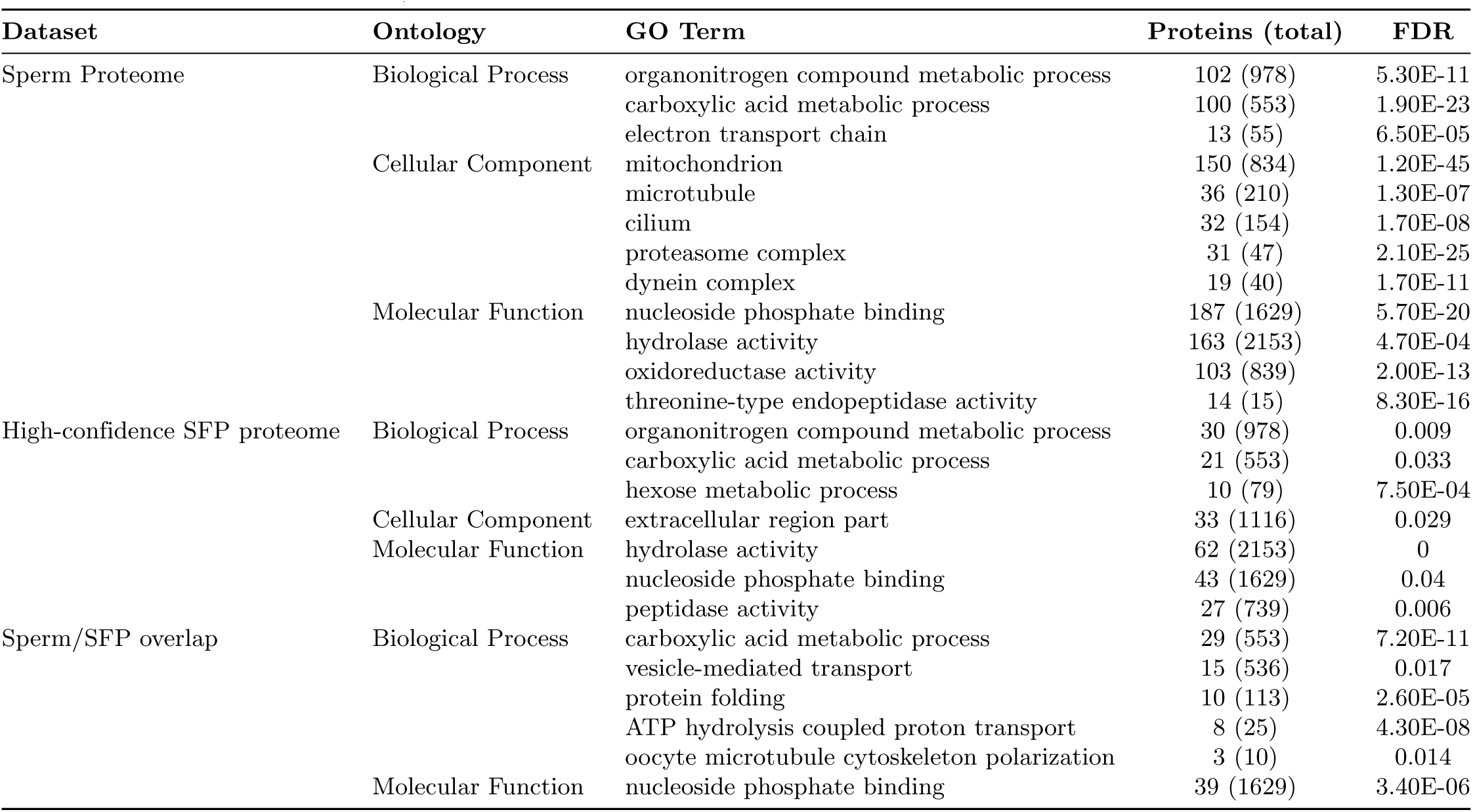
Gene Ontology analysis of sperm proteins, high-confidence SFPs, and 1072 sperm/SFP overlap. List of significant terms is abbreviated to exclude redundancy and to focus on terms discussed in text. 1074 For exhaustive list, see Table S3. FDR; false discovery rate.

In our high-confidence SFP proteome, extracellular structure was significantly enriched amongst cellular component categories, further supporting the accuracy of our SFP identification. Over represented biological processes include proteolysis and both carbohydrate and amide metabolism. Significantly enriched molecular functions include hydrolase activity and peptidase activity. This observation is consistent with the widespread presence of proteolytic enzymes and regulators in SFPs of other insects (reviewed in [64]). The sperm/SFP overlap proteome shares several enriched categories with both the sperm and high-confidence SFP proteomes, including carboxylic acid metabolic processes and nucleoside binding. Several new GO terms emerge in this protein set as well, including ATP hydrolysis coupled proton transport, vesicle-mediated transport, and protein folding (Table 1; Table S3).

### Orthology with sperm proteins and SFPs in other species

We next examined orthology of sperm proteins and SFPs in two different species: *D. melanogaster,* given its well-characterized sperm proteome and SFPs [32,32], and *Ae. albopictus,* which is the closest species to *Ae. aegypti* with characterized SFPs [28]. Orthology was determined between the complete genome of all three species, and then orthologs of *Ae. aegypti* SFPs and sperm proteins also classified as SFPs or sperm proteins in the other species were identified. In the comparison with *Ae. albopictus,* we focus solely on SFPs because a thorough sperm proteome is lacking. Overall, μ87% and μ98% of proteins in the *Ae. aegypti* genome have an ortholog (either as one to-one or one-to-many relationships) in *D. melanogaster* and *Ae. albopictus,* respectively. Among our identified proteins unique to the *Ae. aegypti* sperm proteome, 760 (99%) have an ortholog in *D. melanogaster,* and 451 (59%) of these are also found in the *D. melanogaster* sperm proteome (Table S2; [32,63]). Out of the 280 *Ae. aegypti* SFPs characterized in our study (including 177 high-confidence and 103 sperm/SFP overlap proteins), 275 (98%) have orthologs in *D. melanogaster.* Of these, only 11 have also been identified as *D. melanogaster* SFPs (4%; Table S2). Proteins that contribute to the seminal fluid proteome have therefore diverged extensively during Dipteran evolution. Orthologs were identified in the *Ae. albopictus* genome for 275 (98%) of our SFPs. Of the *Ae. albopictus* SFPs identified to date [28], 86 (43%) were classified as SFPs in our study (Table S2).

### MAG and testis transcriptome characterization and differential expression

We used short-read RNA sequencing to examine gene expression in testes and MAGs to (1) identify transcripts with tissue-biased expression, (2) compare transcript and protein abundance, (3) assess the transfer of male RNAs to females during mating and (4) characterize the effect of mating on MAG gene expression. In both MAG and testes, we find a significant correlation between transcript abundance and protein abundance for high confidence SFPs and sperm proteins (SFPs: R^2^ = 0.57, *F* = 110.4, *p <* 0.001; sperm: *R*^*2*^ = 0.45, *F* = 223.8, *p <* 0.001; Figure 2A-B). As such, variation in transcript abundance explains a substantial amount of protein variation in both of our samples. In total, ˜11,000 and μ7,000 genes had detectable mRNA expression in the testes and MAG, respectively. However, only a subset of these show >2-fold expression bias in testes or MAG compared to other tissues (testes: 2,863; MAGs: 1,485). We examined the association between tissue-biased mRNA expression and differential protein abundance for proteins that were detected in both ejaculate and sperm samples. These results demonstrate that proteins with significant protein abundance differences between sperm and ejaculate samples also tend to show >2-fold tissue-biased mRNA expression (Figure 2C and 2D), further supporting our SFP classification criteria (see above). However, we note that this relationship is less faithful for lower abundance proteins.

**Figure 2.**
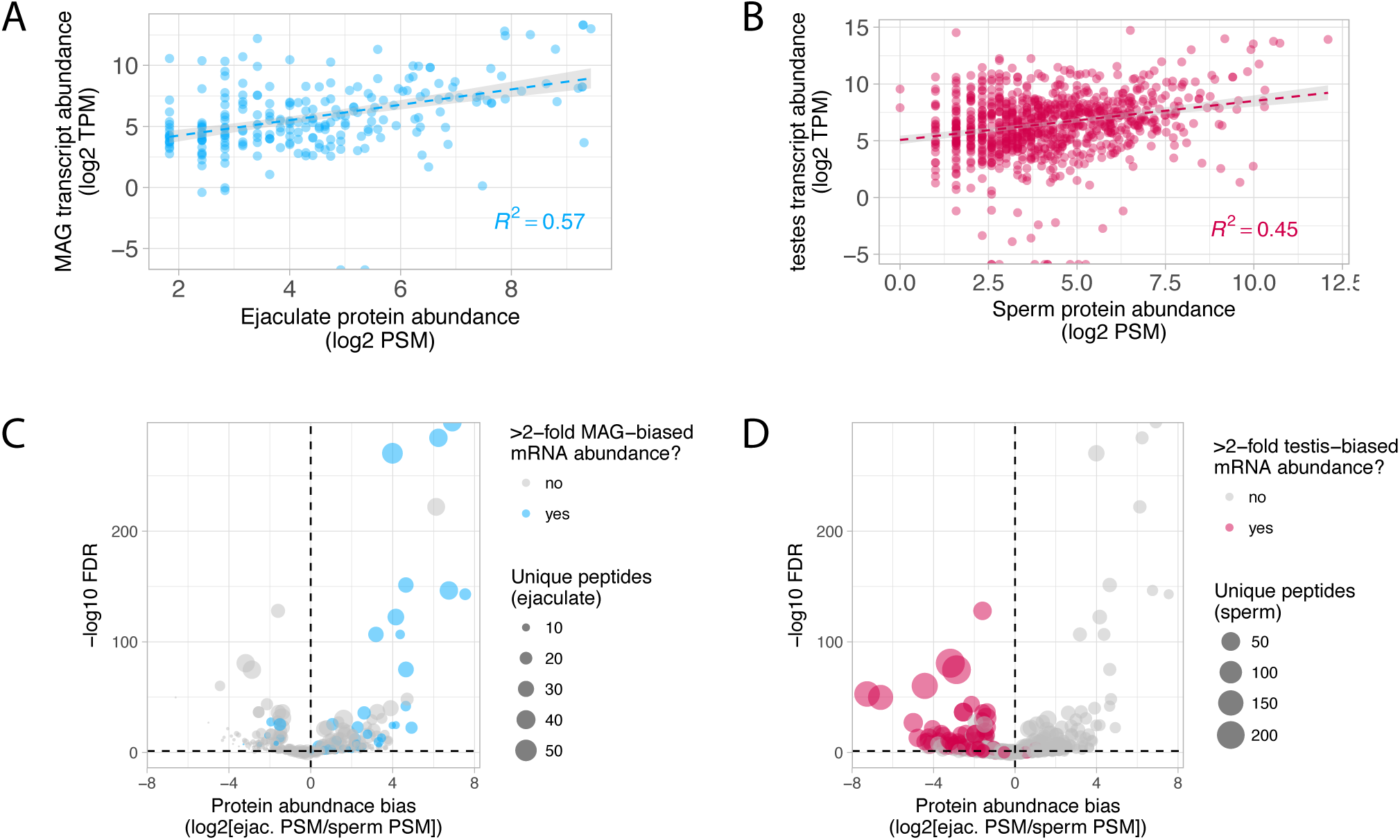
Protein and mRNA abundance relationships. **(A)** and **(B)** Scatterplots of normalized mRNA and protein abundance in MAG/ejaculate (A) and testes/sperm (B). The line and gray shading represent a linear model fit and its 95% confidence interval, respectively. Correlation coefficients are indicated. **(C)** and **(D)** Volcano plots of protein abundance differences for all proteins detected in both the ejaculate and sperm samples. Proteins that show >2-fold mRNA expression-bias in MAG or testes tissue are indicated in blue (MAG-biased) or red (testes-biased), and the size of each point corresponds to the number of unique peptides detected for each protein.

Because males regenerate seminal fluid over the course of 48 h after depleting their reserves by repeated insemination [46], we reasoned that MAG transcriptional regulation after mating might inform our understanding of pathways required to restore depleted SFPs. Differential expression analysis of virgin and mated MAGs’ transcriptomes revealed a significant bias towards gene upregulation in mated males, with 320 transcripts that are upregulated and 126 that are downregulated after mating (binomial test; *p <* 0.001; Figure 3A). In contrast to downregulated transcripts—which are not enriched for any functional category—upregulated transcripts are enriched for several GO categories, many of which are consistent with this tissue’s primary function of producing secreted proteins (Table 2). For example, amino acid metabolism and aminoacyl-tRNA ligase activity are enriched. Enrichment of proteins involved in ubiquitin-dependent protein catabolism and those making up components of the proteasome suggests that there may be extensive protein recycling as seminal fluid is regenerated. Furthermore, the over representation of proteins associated with endoplasmic reticulum targeting and signal peptide processing is consistent with the vesicle-mediated apocrine secretion by which some of mosquito seminal fluid is reported to be produced [60]. We also note that a variety of immune-related genes are upregulated in MAGs after mating, including a family of Defensin antimicrobial genes (Figure 3A). Lastly, a similar pattern was observed among our characterized SFPs, of which 19 were upregulated after mating and only five that decreased in abundance (binomial test; *p* < 0.001; Figure 3B). Interestingly, upregulated SFPs included six different cytoplasmic tRNA ligases and a catalytic subunit of a signal peptidase responsible for post-translation removal of secretion signal peptides, consistent with the importance of protein production and secretion in the mated MAG (see Discussion).

**Table 2.**
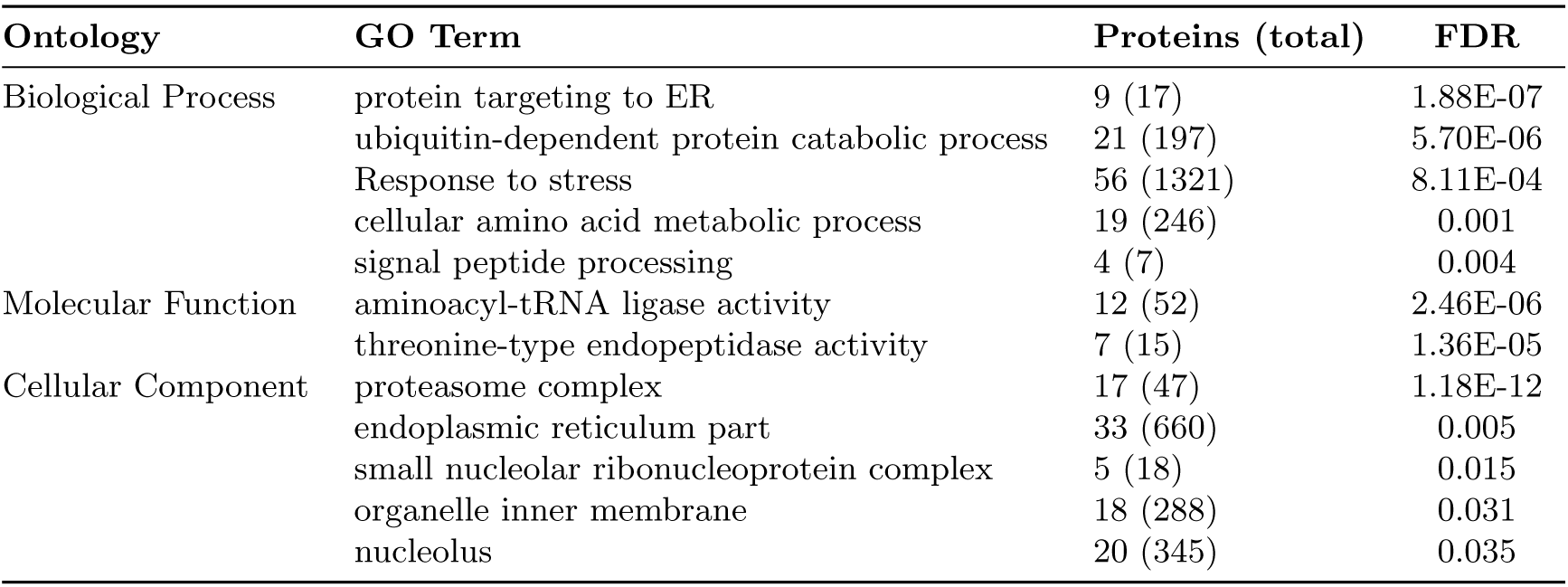
Gene Ontology analysis of upregulated genes in MAGs after mating.

**Figure 3.**
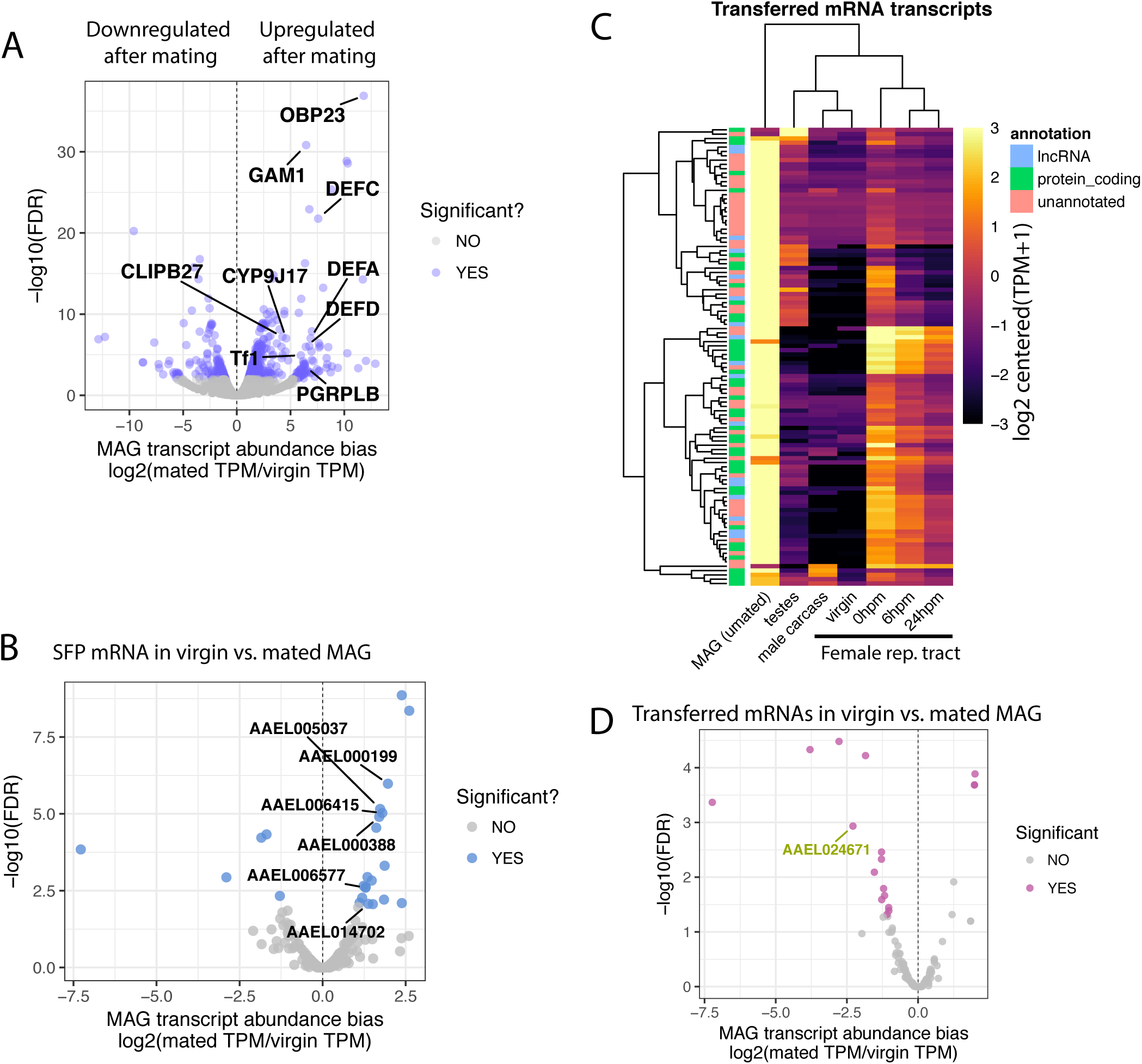
Differential expression between virgin and mated MAGs and abundance of transferred mRNAs. **(A.)**Volcano plot of the 446 differentially abundant RNAs between virgin and mated MAGs. Immune-related genes upregulated after mating are highlighted. **(B)** Volcano plot displaying differential expression of SFP-encoding RNAs between virgin and mated MAGs. tRNA ligase transcripts that are upregulated after mating are highlighted. **(C)** Heatmap of the 106 transferred RNAs in the MAG and testes samples, female reproductive tract samples [49], and a gonadectomized male carcass sample [48]. The annotation classification of each transcript is indicated on the left. **(D)** Volcano plot of the 106 putatively transferred mRNAs and their differential expression status between virgin and mated MAGs. A single lncRNA that shows reduced abundance after mating is indicated.males compared to MAGs of virgins, including 13 transferred transcripts that were 466 significantly down-regulated after mating between virgin and mated MAGs (Figure 3C); 467this pattern may be due to their transfer without replenishment by the MAG at the time of dissection.

### Paternal mRNA transfer during mating

Previously, we have shown that males transfer mRNA to females in the ejaculate [49]. Using the newly annotated genome [38], we re-analyzed data from those experiments and identified 106 transcripts, including 41 protein coding genes and 17 long non-coding RNAs, that are putatively transferred to females. Identification was based on >2-fold increase in transcript abundance in females immediately after mating, followed by a subsequent decrease in transcript abundance. The remainder of the identified transcripts (48 of 106) are currently unannotated. Using the transcriptomic data in the current study, we determined that the vast majority of these transcripts have MAG-biased expression, with only two transcripts exhibiting testis-biased expression (Figure 3C). The MAG therefore appears to be a primary source of RNA transferred to females in the ejaculate. In total, 27 proteins encoded by transferred mRNA transcripts were identified in our proteomes, with 22 in the high-confidence SFP proteome, three in the sperm/SFP overlap proteome, and five that were in the sperm proteome. Interestingly, the putatively transferred transcripts whose products were present in our seminal fluid proteome encode highly abundant proteins that were on average six times more plentiful than the remainder of the seminal fluid proteins. Lastly, transferred transcripts exhibited a general trend towards down-regulation in the MAGs of mated

### SFPs are enriched on chromosome 1

Sex chromosomes present exclusively in males, such as the mammalian and insect Y chromosome, are highly enriched for genes with male-biased function, including many critical to spermatogenesis and sperm function [32,65,66]. Although *Ae. aegypti* lack heteromorphic sex chromosomes, chromosome 1 harbors a region of robust linkage disequilibrium surrounding the recently characterized male sex determination locus, *Nix* [55,67]. To assess the enrichment of male-biased genes on this chromosome we examined the physical distribution of SFPs, sperm proteins, and MAG/testes-biased genes by chromosome. This revealed a μ1.5-fold enrichment of SFPs on chromosome 1 (62 observed, 39 expected; x^2^ = 12.4, *p <* 0.001) and μ1.2-fold enrichment of sperm proteins on chromosome 2 (377 observed, 316 expected; x^2^ = 11.6, *p <* 0.001; Figure 4). Despite the strong enrichment of SFPs on chromosome 1, we did not observe an over-representation of SFPs in the tightly linked region in the vicinity of the male determining locus, *Nix* (15 observed, 16 expected; x^2^ = 0.08, *p* = 0.8). Together, these results suggest that chromosome 1 does exhibit aspects of male-biased functional composition, but that this signature is not readily detectable in the linked region that harbors the male sex-determining factor *Nix*. Non-uniform physical distribution of sperm protein across autosomes has also been observed in *Drosophila* [63]. Although the functional significance of this remains to be determined, it is possible that clustering facilitates co-expression during spermatogenesis, which is characterized by progressive genome silencing during the histone-to-protamine repackaging transition.

**Figure 4.**
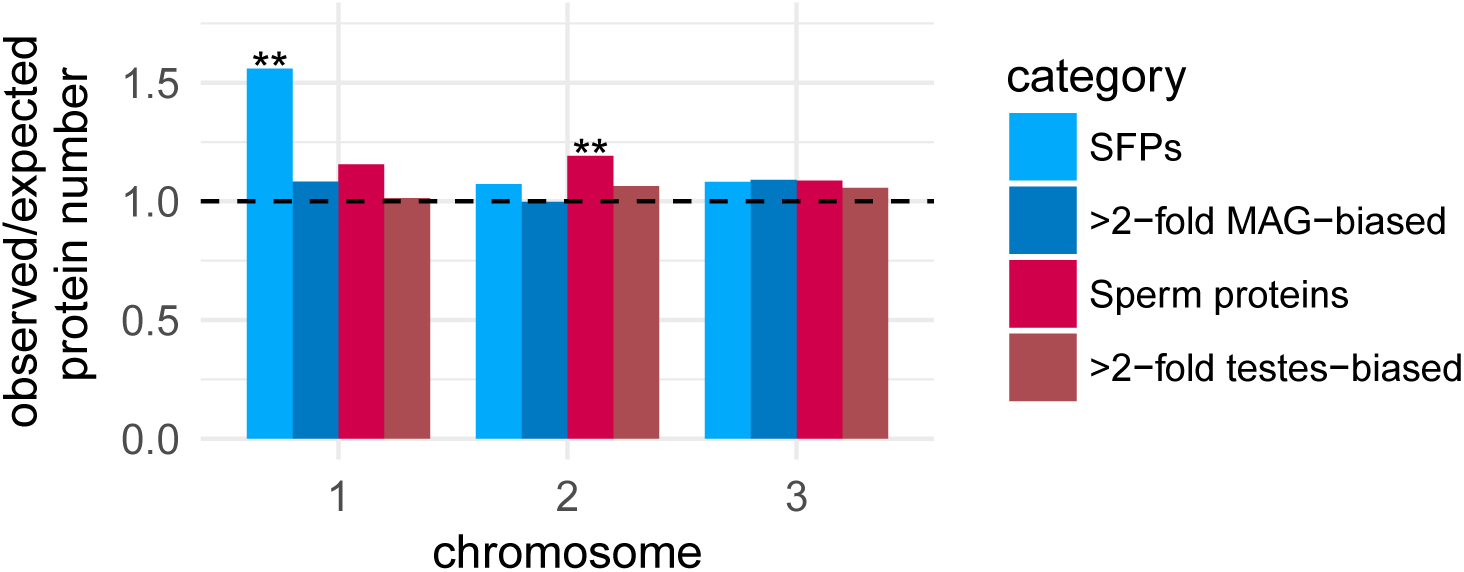
Chromosomal distribution of genes encoding SFPs, sperm proteins, or transcripts with MAG biased expression or testis biased expression.

The *y*-axis represents the ratio of observed/expected number of genes for each of the chromosomes in *Ae. aegypti,* and the dashed line represents the expectation under no enrichment/depletion. Asterisks indicate significant enrichment (x_2_; **p < 0.01).

## Discussion

A rapidly expanding body of evidence supports the critical roles of seminal fluid proteins (SFPs) in a wide array of reproductive phenotypes (reviewed in [29]). Although this has been most extensively investigated in *Drosophila*, seminal peptides are also associated with post-mating behavioral and physiological responses in mosquitoes such as *Ae. aegypti* [15,16,18]. The primary goals of this study were to comprehensively catalog male proteins transferred to *Ae. aegypti* females during insemination and establish a reliable methodology for delineating between sperm proteins and SFPs. To accomplish this, we (1) conducted an in-depth proteomic characterization of sperm, (2) utilized a whole-female labelling approach to identify unlabelled male proteins transferred by the male during insemination and (3) characterized the transcriptomes of the testis and male accessory gland (MAG). Importantly, we note that the whole-female labelling approach has been employed previously in *Ae. aegypti* but the assignment of proteins as SFPs was limited by the lack of information regarding proteins found in sperm. Thus, distinctions between sperm proteins and SFPs were previously difficult to achieve. It is also noteworthy that advances in MS/MS sensitivity and accuracy have resulted in far greater power of detection in our study, and our analysis has also benefited tremendously from the recent resequencing and reannotation of the *Ae. aegypti* genome [38]. Our proteomic characterization resulted in a nearly four-fold expansion of the current *Ae. aegypti* seminal fluid proteome, and an eight-fold expansion of identified proteins in sperm. Together, this *Ae. aegypti* “ejaculatome” provides a foundation for future molecular studies of mosquito reproduction and associated applications to control mosquito populations (see below).

### Proteome characteristics independently validate identification

Our work differs from previous SFP characterization studies [27,28,57] in that our classification was supported by a detailed knowledge of sperm proteome composition. Nonetheless, several independent validation approaches were helpful in assessing the quality of our proteomic characterization. For example, we quantified the proportion of proteins with predicted secretion signals and analyzed transcriptome profiles in testes and MAGs. As would be predicted, SFPs identified in this study possessed a significantly higher proportion of predicted secretion signals than sperm proteins and were, on average, highly specific or biased towards expression in the MAG. Additionally, analysis of the functional composition of our proteomes revealed that they were closely aligned with the results of previous sperm [32-34,63] and SFP studies in insects [30,57]. For example, our expanded sperm proteome was highly enriched for proteins related to flagellar structure, including microtubules, dynein complexes, and ciliar components, and proteins likely associated with the mitochondrial derivatives, which are a predominant structure in mosquito sperm [68,69] and that of other insects. Consistent with what has been described by Sirot *et al.* [27], as well as in other insects (reviewed in [29,30,57]) and humans [70], proteases were highly enriched amongst our high-confidence SFPs, supporting the likely accuracy of our expanded characterization (reviewed in [64]). The observed enrichment of vesicle-mediated transport proteins is also consistent with the fact that mosquito seminal fluid is in part produced by apocrine secretion [60]. Additionally, exosomes and other vesicles are believed to play a role in a variety of post-insemination cellular interactions. For example, vesicles transferred in *Drosophila* seminal fluid have been reported to fuse with sperm and interact with the female reproductive tract [71], exosomes of the mouse epididymis have recently been implicated in the control of sperm RNA stores [72], and the abundance of exosome markers in avian SFPs has led to speculation about vesicle-mediated mechanisms in post-testicular sperm maturation [73]. Therefore, the accuracy of our expanded proteomic characterization of sperm and SFP proteomes is corroborated by several independent lines of evidence.

It is important to note that, despite the application of stringent proteomic thresholds, some proteins could not be definitively assigned as either sperm protein or SFP. Previous studies in *Drosophila* and Lepidoptera have consistently identified known SFPs (such as Acp36DE) at appreciable abundance levels in sperm that have yet to be combined with MAG secretions [32,62,63]. Our identification of a relatively large protein set that is highly MAG-biased in expression but also present in sperm further suggests that the incorporation of “SFPs” during testicular sperm maturation occurs and is worthy of additional functional investigation. Although *Drosophila* expression profiles in the testis and accessory gland are quite distinct, many SFPs exhibit low levels of co-expression in the testis (Dorus, unpublished data). Our transcriptomic analyses here further support such patterns of co-expression. As such, dichotomous distinctions between sperm proteins and SFPs may be an oversimplification of a more nuanced relationship between these reproductive systems. We acknowledge this uncertainty in our classification of MAG-biased proteins that were also identified in our sperm proteome.

We also note that despite our expanded proteomic coverage, several proteins that we anticipated to be identified were absent. The most notable case was Head Peptide-1, a seminal fluid peptide which has been shown to be transferred in the ejaculate [74] and has been reported to induce short term monogamy in the female after mating [15]. Head Peptide-1, as is the case for many SFPs, undergoes extensive post-translational modification and may therefore be challenging to identify bioinformatically without a priori knowledge of the biochemical composition of the proteolytic products (such as in the case of the well-studied *Drosophila* Sex Peptide; [75]). Another example was adipokinetic hormone (AAEL011996), which did not meet our two unique peptide inclusion threshold, although we did identify five copies of one peptide from its precursor protein that was also identified in *Ae. albopictus* seminal fluid [28]. This protein has been postulated to contribute to sperm protection from oxidative stress [76] and the regulation of feeding behavior [77] in other insects. We suggest that the complexity of proteolytic pathways, governed both by male and female interacting proteins, is a major barrier in the use of shotgun proteomics to study SFP identity and function in the female reproductive tract. Future investigations would likely benefit from the inclusion of a targeted proteomic approach (reviewed in [78]). Such approaches require an a priori list of candidate peptides; in *Ae. aegypti,* the neuropeptides and protein hormones catalogued by Predel *et al.* [79] represent a useful pool of potentially important molecules.

### Evolution of male reproductive proteomes

Male reproductive proteins, including SFPs, are consistently among the fastest evolving classes of protein (reviewed in [80]). Although initially a goal of our study, conducting a robust analysis of the molecular evolution of protein identified in this study was limited by the availability of genomic resources appropriate for both inter- and intraspecific tests of positive selection. Obtaining high quality genomic data for different populations of *Ae. aegypti* has proven difficult, given the genome’s repetitive nature [38,81]. Furthermore, this mosquito’s ability to move globally as diapausing eggs has allowed for frequent mixing and a complex population structure [82,83]. The development of appropriate population level genetic data for the analysis of recent selective sweeps should be a priority in *Ae. aegypti,* as it has been in Anopheles gambiae [84,85]. Furthermore, given the extent of molecular divergence between *Ae. aegypti* and *Ae. albopictus* [28], the development of genomic resources for a more closely related outgroup to *Ae. aegypti* will assist in understanding evolutionary patterns at the gene level. Despite these limitations, our analysis of orthology did reveal that the suite of proteins contributing to seminal fluid, but not sperm, has diverged substantially from other Dipterans. Although sperm proteins and SFPs possess levels of orthology to the *Drosophila* genome that are comparable to the genome as a whole, only 59% and 4% of orthology was observed when comparing the *Ae. aegypti* sperm and SFP proteomes (respectively) with those of *Drosophila* [32,57]. While some of this disparity may be attributed to differences in overall proteome size and coverage, such a stark contrast is nonetheless compelling evidence of tissue-specific evolutionary patterns. Orthology between *Ae. aegypti* SFPs and *Ae. albopictus* SFPs [28], while more extensive (43%), was still comparatively low compared to orthology between the sperm proteomes of *Ae. aegypti* and *D. melanogaster*—two distantly related Dipterans. These results suggest a process of “turn-over” in seminal fluid proteomes, whereby overall protein composition diverges rapidly even when there is evidence for conservation with regard to overarching molecular functions represented in seminal fluid. For example, a priori expectations about Gene Ontology enrichment were met for both *Ae. aegypti* sperm *(e.g.,* cilium and mitochondrial proteins) and SFPs (extracellular localization and hydrolase activity), despite overall SFP divergence. SFPs are a pronounced target of selection and have been discussed as a driver of sexual conflict (reviewed in [86]), and thus they are expected to rapidly diverge. By contrast, we note that strong conservation of sperm proteins exists across distant taxa, with different insect orders displaying 25% orthology between sperm proteomes [34], and even *D. melanogaster* and mammals with 20% sperm proteome orthology [32]. The overall lack of conservation in seminal fluid proteomes makes comparing the roles of specific SFPs across species difficult, but conserved molecular functions amongst SFPs will nevertheless allow the wealth of knowledge in *Drosophila* to be leveraged towards an understanding of SFP function in non-model insects.

Unlike other mosquitoes with heteromorphic sex chromosomes, Culicine mosquitoes *(e.g., Aedes* and *Culex*) harbor male determining loci on undifferentiated, homomorphic sex chromosomes [87]. Theory predicts the evolution of heteromorphic sex chromosomes following the acquisition of a sex determining locus, suppression of recombination, and expansion of the non-recombining region. It remains unclear why homomorphic sex chromosomes appear to be retained in some taxa [88,89]. One proposed mechanism to mediate the selective effect of sexually antagonistic alleles on the promotion of recombination suppression is the establishment of efficient sex-b iased expression [90]. Although previously lacking, the significant enrichment of SFPs on chromosome 1 is the first evidence in support of this hypothesis in *Ae. aegypti*. This trend was restricted to SFPs and was not observed for genes solely over-expressed in the MAG or testis. It is intriguing to speculate that this distinction between SFPs and other male reproductive genes might be due to the prevalence (and selective strength) of sexually antagonistic alleles specifically amongst SFPs, which may favor their localization on chromosome 1. This is consistent with their putative role as drivers of sexual conflict (reviewed in 86), including the mediation of female post-mating responses such as sexual receptivity and longevity [14,18].

### Functional relevance of abundant sperm proteins and SFPs

Our sperm and SFP proteomes exhibited skewed abundance distribution with the top ten most abundant proteins comprising 25% and 17% of protein composition in sperm and SFPs, respectively. Interestingly, the most abundant sperm protein, cytosol aminopeptidase (AAEL006975), accounted for more than 7.4% of all protein and two other cytosol aminopeptidases were in the top ten most abundant proteins (AAEL000108, AAEL023987). These three proteins are orthologs of the eight sperm-leucyl aminopeptidases (S-LAPs) in *Drosophila* with similar expression patterns, including μ1000-fold higher expression in testes than in MAG, μ50 times more transcript in whole male carcasses than gonadectomized carcasses [91], and upregulation during later stages of spermatogenesis [48]. S-LAP orthologs constitute a significant proportion of the protein composition of *Drosophila* [92] and Lepidoptera [34,62]. Little is known about the specific function of S-LAPs, although it has been postulated that may serve a structural function given the inferred loss of enzymatic capacity of several S-LAPs during *Drosophila* evolution [92]. Additionally, a Y-linked S-LAP in D. pseudoobscura has been implicated in a cryptic meiotic drive system, where suppression of this locus results in aberrant spermatogenesis and a higher proportion of X-bearing sperm [93]. It will be of great interest to establish the specific function of these proteins in mosquito sperm, given their high abundance and expression patterns during spermiogenesis. Furthermore, the proteins and transcripts involved in spermatogenesis described in this study may assist in the identification of other genes involved in meiotic drive systems (reviewed in [94]), which have been proposed as potential genetic means to reduce wild populations through the induction of sex ratio biases [95].

Although no SFP was as abundant as cytosol aminopeptidase in sperm, the top ten most abundant proteins ranged from 1.2 - 2.6% of the protein in our ejaculate sample. L-asparaginase (AAEL002796) was the most abundant SFP (61% more abundant than the next protein) and the tenth most abundant mRNA transcript in the MAG out of over 11,000 transcripts. While the relevance of the abundance of this enzyme is currently unclear, it may relate to several other notable observations. First, transcript AAEL020035, whose protein product is comprised of μ60% asparagine residues, is the single most abundant MAG transcript and was also, by far, the most abundant putatively transferred transcript. (We did not identify AAEL020035 in our SFP proteome but note that it results in few identifiable peptides because of its extreme amino acid composition). Second, asparagine tRNA-ligase was two times more abundant in seminal fluid than any other tRNA-ligase. Third, asparagine tRNA-ligase was upregulated in the MAG after mating and was the most abundant tRNA-ligase transcript. Together, these suggest that MAGs are well-equipped to produce ample protein with an asparagine amino acid bias. Finally, two other enzymes, aspartate transaminase (AAEL002399) and citrate synthase, are abundantly present in seminal fluid and could convert aspartate produced by asparaginase to oxaloacetate and citric acid, respectively. While it is premature to draw any firm conclusions based on these observations alone, it is intriguing to speculate that the SFP proteome has the capacity to conduct gluconeogenesis (of asparagine and potentially other amino acids) and that this may feed into to the citric acid cycle. The citric acid cycle is believed to be functional in mammalian sperm (reviewed in [96]) and many citric acid cycle enzymes are present in our *Ae. aegypti* sperm proteome.

The most abundant seminal fluid proteins also exhibited a strong enrichment for function in protein cleavage. Proteins with protease, dipeptidase, and aminopeptidase activity represent 12 of the top 28 most abundant proteins present in the seminal fluid proteome. Proteolytic functions have been described previously in the seminal fluid of *Ae. aegypti* [27], *Ae. albopictus* [28], *Cx. quinquefasciatus* [97], and several non-mosquito taxa [21,22](reviewed in [64,98]), and are a common function of many insects’ seminal fluid. Based on studies in other insects, functions of these enzymes may include the activation of sperm motility or the cleavage of propeptides into their active forms [99]. Our seminal fluid proteome also contains abundant enzymes that catabolize smaller substrates, such as amino acids and carbohydrates. Taken together, the enzymatic cocktail in seminal fluid may be well equipped to break down many of the molecules they contain. Seminal fluid proteins were also enriched for proteins involved in maintaining proton and redox homeostasis. We identified several proteins contributing to V-type proton ATPases, which use ATP to regulate pH via proton transport. Maintaining an optimal pH in seminal fluid may allow for efficient sperm motility (reviewed in [100]). Regulating pH may also create an ideal environment in which enzymatic reactions occur. Seminal fluid also contained several proteins that function to neutralize free radicals, such as catalase (AAEL013407), peroxidase (AAEL013171), and several dehydrogenases. Regulating the physiochemical environment in seminal fluid is likely critical for the function and protection of sperm prior to their storage in the female’s long term storage organs (spermathecae).

### Ejaculate RNAs Transferred to Females

There has been much conjecture about the importance of spermtazoal RNA to fertility [101] and recent work has confirmed that the regulation of sperm ncRNA stores in the mammalian epididymis is necessary for proper embryogenesis [102]. Little is known about the function of spermatozoal RNAs in insects, although they have been demonstrated to have substantial functional coherence, including an overwhelming enrichment of loci involved in translation [103]. New data in this study allowed us to probe for patterns in previously described transcripts that are putatively transferred to females during mating. A total of 106 transcripts were identified, including both coding and non-coding transcripts, and a majority of these exhibit high levels of expression in the MAG. Based on our SFP proteome, most of the protein coding transcripts are translated at high levels. Their high expression in the MAG suggests that they may simply hitchhike into seminal fluid with other secreted molecules. Alternatively, as has been demonstrated in *Drosophila,* they could be transferred in intact MAG cells [61], or via vesicles derived from the MAG [71]. Interestingly these vesicles, which may carry RNA cargo including miRNAs, fuse with sperm and have the capacity to interact with the female reproductive tract. Some male-derived transcripts are detectable in the female for up to 24 h post-mating [46], and it has been postulated that they could be used by females in some capacity [104]. In *Ae. aegypti,* both vesicles and RNAs are transferred in the ejaculate to the female, but their fate and function have not been investigated. Whether they impact the female or her future offspring is an intriguing, and potentially important, line of future investigation.

### Mosquito control and future directions

Understanding the molecular architecture of *Ae. aegypti* reproduction holds great potential for vector control strategies. Mosquito reproduction is an ideal control target to reduce vector populations and the burden of disease transmission. The most direct application of this study will be the identification of modulators of female reproductive behavior. Mosquito SFPs induce behavioral responses that prevent female remating [14,16,17], including short term mating refractory behavior [15]. To date, the molecule(s) responsible for long term refractoriness has yet to be identified. Given the strength and duration of responses to low SFP “doses” [14], identification of the responsible proteins will provide powerful tools for manipulating female reproduction in a species-specific manner. In addition, such knowledge may provide a molecular metric by which the quality of males in modified mosquito release strategies (such as those employing sterile or Wolbachia-infected males; reviewed in [105]) may be monitored and optimized. Functional analysis of specific sperm proteins and SFPs may yield insights into processes such as sperm motility and activation [21-23], sperm storage [106], and sperm-egg recognition [107]. Very few studies have explored these processes in *Ae. aegypti* (reviewed in [108]). A mechanistic understanding of complex post-copulatory male-by-female interactions is critical to genetically modified mosquito release strategies that manipulate reproduction. Our detailed characterization of the male contributions to these interactions should serve as the foundation for the design and improvement of vector control strategies that limit the transmission of arboviruses that cause serious human illness and mortality.

## Supporting Information

**Figure S1**

**One-dimension SDS-PAGE gel separation of samples.** Sperm (A) and ejaculate (B) protein fractions (1 - 6) analyzed by LC-MS/MS. Biological replicates were run in parallel on 10% bis-tris SDS-PAGE gel with MOPS running buffer and stained with colloidal Coomassie. V, virgin bursae; M, mated bursae. Labeling efficiency of females was verified through the analysis of fraction 5 from virgin females.

**Table S1**

**Database of all proteins identified by tandem mass spectrometry in this study.** Classification of each protein is based on criteria in text. Proteomic (probability of identification, percent coverage, total PSMs, and unique PSMs) and transcriptomic (TPM in virgin MAG, mated MAG, and testes) data are included, as well as whether each protein includes a signal peptide, and how each protein was previously classified by Sirot *et al.* [27].

**Table S2**

**Full GO analysis for high confidence SFPs, sperm/SFP overlap, and sperm proteins.** CC, cellular component; BP, biological process; MF, molecular function.

## Acknowledgments

We thank Sylvie Pitcher, Sheng Zhang, Jen Grenier, Peter Schweitzer, and the staff at the Cornell Biotechnology Resource Center for technical support, and Laura Sirot for experimental guidance and feedback. This study was supported by *NIH/NIAID grant R01AI095491 to MFW and LCH, NIH/NICHD grant R21HD088910 to SD and MFW, a Cornell Graduate School fellowship to ECD, and a Cornell Entomology Department Griswold grant to ECD and LCH. YHAB was supported by NIH/NICHD grant R01HD059060 to MFW and Andrew G. Clark. RNA-seq data and mass spectrometry data were made possible by NIH grants 1S10OD010693-01 and 1S10OD017992-01, respectively.

